# Reconciling biodiversity conservation, food production and farmers’ demand

**DOI:** 10.1101/485607

**Authors:** Daniel Montoya, Sabrina Gaba, Claire de Mazancourt, Vincent Bretagnolle, Michel Loreau

## Abstract

Agricultural management should consider multiple services and stakeholders. Yet, it remains unclear how to guarantee the provision of ecosystem services that reaches stakeholders’ demands, especially considering the observed biodiversity decline and the current global change predictions that may affect food security. Here, we use a model to examine how landscape composition – fraction of semi-natural habitat (SNH) – affects biodiversity and crop production services in intensively-managed agricultural systems. We analyse three groups of stakeholders assumed to value different ecosystem services most – individual farmers (crop yield per area), agricultural unions (landscape production) and conservationists (biodiversity). We find that trade-offs among stakeholders’ demands strongly depend on the degree of pollination dependence of crops, the strength of environmental and demographic stochasticity, and the relative amount of an ecosystem service demanded by each stakeholder, i.e. function thresholds. Intermediate amounts of SNH can allow for the delivery of relatively high levels of the three ecosystem services. Our analysis further suggests that the current levels of SNH protection lie below these intermediate amounts of SNH in intensively-managed agricultural landscapes. Given the worldwide trends in agriculture and global change, these results suggest ways of managing landscapes to reconcile the demands of several actors and ensure for biodiversity conservation and food production.

## 1. INTRODUCTION

Natural ecosystems provide a wide range of benefits to people. The concept of ecosystem services (Ehrlich and Ehrlich, 1981) was originally proposed to draw attention to these benefits and raise awareness about the importance of biodiversity and its conservation. More recently, the ecosystem service metaphor has incorporated two additional elements. First, ecosystems provide multiple services simultaneously to humans (provisioning, supporting, regulating and cultural), a situation called multifunctionality (Manning et al., 2018). This has led to a growing consensus that landscape design and management should target a range of functions (Wilson, 2007; Shellhorn et al., 2008; Birkhofer et al., 2015; Landis, 2017). Second, the ecosystem service concept is a complex construct that comprises both supply (ecosystems) and demand (stakeholders) components (Burkhard et al., 2012; Palacios-Agundez et al., 2015; Yahdjian et al., 2015). Human use of ecosystem services thus depends on the capacity of ecosystems to supply these services as well as on the social demand for them.

Agricultural landscapes include a great variety of ecosystem services and stakeholders, and they globally represent about 40% of the total terrestrial surface of our planet (Foley et al., 2011). As such, agricultural landscapes are among the most interesting systems to analyze the balance between the supply and demand of various ecosystem services. The particularity of most modern agricultural landscapes is that the provision of a single service, i.e. crop production, is intensified by means of land conversion, mechanical work and the use of agrochemicals (pesticides, fertilizers). But this generates negative indirect effects, as land use favors crop production at the cost of other services that in turn influence crop production (Nelson et al., 2009; Allan et al., 2015; Sutter and Albrecht, 2016). The need to incorporate various ecosystem services into management decisions is thus essential, especially considering that agricultural intensification is no longer enhancing the yields of many major crops worldwide (Ray et al., 2012), and the response of crops to the use of pesticides is saturating (Gaba et al., 2016; Lechenet et al., 2017). This has been acknowledged by the 2017 UN Sustainable Development Goals, which assert that the future of intensive farming systems requires management strategies that achieve food security and sustainable agriculture.

The supply and demand components of ecosystem services in agricultural systems are multifaceted because of the multiple ecological processes involved on the supply side and the multiple stakeholders that benefit from them on the demand side. On the supply side, trade-offs between ecosystem services seem to be common (Bennett et al., 2009; Nelson et al., 2009; Raudsepp-Hearne et al., 2010; Allan et al., 2015; Ruijs et al., 2013; Divinsky et al., 2017; Villasante et al., 2016). Intensive agricultural management, for example, may lead to high crop yields, but intensively managed fields often have simplified communities and hence low levels of other ecosystem services such as biological control by natural enemies and pollination (Médiène et al., 2011). Such trade-offs depend on various factors, including landscape composition (crop land *vs* semi-natural land) and the degree of pollination dependence of crops (Aizen et al., 2009; Garibaldi et al., 2014, 2011a, 2011b; Deguines et al., 2014; Montoya et al., 2018). Together these factors lead to scenarios where either different ecosystem services can be reconciled or, conversely, the provision of several ecosystem services is incompatible. Evaluating landscape performance when multiple interacting factors act simultaneously is thus a challenge, especially considering questions such as which combination of drivers is able to reconcile crop production and biodiversity conservation.

Human demand is the other side of the ecosystem-service equation, and comprises different agents or stakeholders. These stakeholders vary in both their demand for and valuation of different ecosystems services, and have specific perspectives about how to efficiently manage agricultural landscapes. For example, agricultural unions or cooperatives often aim at maximizing crop production at the regional level, i.e. landscape production, for food security purposes. Farmers, on the other hand, may have different perceptions of ecosystem services (Teixeira et al., 2018); yet, they are mainly interested in maximizing crop yield per area of their cultivated land as this is mostly related to their profitability. Conservationists (NGO’s, wildlife-friendly organizations) defend that the preservation of the remaining biodiversity within the agricultural landscape should be one main goal of agricultural policies. The existence of ecosystem service trade-offs on the supply side implies that any stakeholder’s demand does not necessarily maximize multifunctionality of agricultural landscapes, because each stakeholder may prioritize one ecosystem service or set of services over others. Considering various stakeholders is thus required to identify potential trade-offs and balance multiple, often conflicting, demands for ecosystems services (Baro et al., 2017; Turkelboom et al., 2018).

Here, we use a model of biodiversity and crop production in intensively-managed agricultural landscapes to derive landscape management solutions for different stakeholders’ demands. We analyze three main groups of stakeholders – individual farmers, agricultural unions and conservationists – and identify three ecosystem services they value most – crop yield per unit of agricultural area, crop production at the landscape scale, and biodiversity, respectively. Using information given by ecosystem service trade-offs, we determine the best landscape composition, defined as the fraction of semi-natural habitat within the agricultural landscape, that corresponds to each stakeholder’s demand, and how this affects the provision of the other ecosystem services. We also define conditions under which stakeholder demands are compatible or not, e.g. when food production and biodiversity conservation are positively correlated. Global change can have important effects on the delivery of ecosystem services, and acts interactively with land use modifications (Shoyama & Yamagata 2014; Ding et al., 2016; Martinez-Harms et al., 2017). Besides, food stability is considered to be one of the major challenges of food security that is missing in ecosystem service research (Cruz-Garcia et al., 2016) and, therefore, considering variations in ecosystem services due to stochasticity is key to assess both ecosystem service supply and demand components. We investigate how changes in stochasticity (expected under global change predictions) and the degree of pollination dependence of crops may affect the best landscape composition for each stakeholder. Specifically, our work has three objectives: (i) to determine the best landscape compositions that correspond to different stakeholders’ demands, (ii) to investigate the effects of a given stakeholder’s demand on other ecosystem services, and (iii) to establish the best landscape composition for ecosystem multifunctionality, i.e. a social average scenario that targets the highest provision of the three ecosystem services beyond any single stakeholder’s demand. Finally, we confront the model outputs with current policies and discuss their efficiency to promote multifunctional agricultural landscapes. Our approach is unique in its attempt to achieve some balance and/or maximize the provision of ecosystem services and stakeholder demands for different crop types (pollination dependence of crops) and environmental and demographic stochasticity scenarios.

## 2. MATERIALS and METHODS

### 2.1. Model description

We extend a model of biodiversity and crop production in a spatially heterogeneous agricultural landscape that incorporates environmental and demographic stochasticity. Food stability is one of the major challenges of food security that is missing in ecosystem service research (Cruz-Garcia et al., 2016), and our model explicitly includes it through the consideration of environmental and demographic stochasticity (Montoya et al., 2018; Appendix A). The model represents intensively-managed agricultural landscapes, where crop land does not harbour significant levels of biodiversity. Spatial heterogeneity is defined by two types of patches: (i) crop land, which is used to grow annual crops with varying degrees of dependence on animal pollination, and (ii) semi-natural habitat, which shelters biodiversity, including wild plants and pollinators. Crop land and semi-natural habitat are linked by pollinators’ foraging movement, and pollinators are assumed to be generalist central-place foragers that feed on both ‘wild’ plants and crops (Kleijn et al., 2015). The model focuses on wild pollinators as they directly depend on semi-natural habitat for nesting, foraging, and refuge. Space is implicitly considered, so that pollinators can potentially feed on all crops and wild plants present in the agricultural landscape, irrespective of the spatial configuration of the landscape.

The model studies three ecosystem services provided by agricultural landscapes – crop yield per unit of agricultural area, crop production at the landscape scale, and biodiversity, and describes mechanistically the trade-offs between these ecosystem services in intensive agricultural landscapes (Montoya et al., 2018). This model is a useful first approximation to studying the crop production in agroecosystems, as it successfully reproduces empirical observations on the stability of pollination-dependent crop yield (Garibaldi et al., 2011a; Deguines et al., 2014), and it provides rigorous theoretical foundations for previously hypothesized functional relationships between the magnitude of ecosystem services and landscape composition (Braat and ten Brink, 2008). Model dynamics are governed by two key parameters: the degree of pollination dependence of crops, which we vary to represent different crop types (see Appendix A), and environmental and demographic stochasticity. Here, we analyse the effects of these parameters on the best landscape composition associated with each stakeholder’s demand. A complete description of the model, the model equations and the estimation of model parameters is provided in Appendix A.

### 2.2. Stakeholders and management scenarios

We use our model to investigate how different stakeholders’ demands determine the best landscape compositions in agricultural landscapes. Beneficiaries or stakeholders for ecosystem services include individuals, cooperatives, corporations, non-governmental organisations, and the public sector. To illustrate our approach, we analysed three main groups of stakeholders – individual farmers, agricultural unions and conservationists –, and identify three ecosystem services valued by those stakeholders most – crop yield per area of their cultivated land, crop production at the landscape scale (landscape production) and biodiversity (scenarios 1-3). We use this example as a case study to determine the best landscape compositions for each different stakeholder’ demands and how they affect the magnitude of the various ecosystems services. Best landscape composition is defined as the range of fraction of semi-natural habitat within which the targeted level of a given ecosystem service is achieved. Additionally, we define a fourth scenario where no single ecosystem service is prioritized; this scenario targets the highest possible provision of the three ecosystem services described above, i.e. multifunctionality (scenario 4), and it can be viewed as a social average scenario beyond the specific demands of the individual stakeholders. The social average scenario follows the idea that a ‘challenge for the future is to design landscapes that are beneficial for a range of functions’ (Shellhorn et al., 2008).

Stakeholders may demand a minimum amount of ecosystem service provision, e.g. protecting ≥75% of biodiversity or producing ≥80% of crop biomass. This is accounted for in our model by incorporating *function thresholds*. A function threshold is defined as the relative amount of an ecosystem service demanded by each stakeholder, i.e. how much of the maximum possible value of that ecosystem service (given by model simulations) a given stakeholder is willing to accept. The best landscape compositions within the agricultural landscape are likely to change if stakeholders assume higher or lower thresholds of their demanded ecosystem service; thus, considering function thresholds is useful for identifying potential trade-offs and for balancing various demands for services.

### 2.3. Analytical protocol

We follow a four-step process summarized in Figure 1. First, for each fraction of semi-natural habitat, we run model simulations to obtain the frequency distribution for each ecosystem service after 1000 time steps (landscape production is used as an example, Fig. 1A). This is performed for the whole range of semi-natural habitat (0-100%) to obtain the frequency distribution of landscape production as a function of semi-natural habitat (Fig. 1B). The strength of this approach lies in its ability to explore the full probability distribution of ecosystem service values as well as the possibility of setting thresholds values for services (see Carnus et al., 2014, for the justification of this approach). Absolute thresholds for ecosystem services are generally lacking, so we look at the best landscape compositions that provide a given proportion of the maximum ecosystem service value along the range of semi-natural habitat (Appendix B provides an alternative application of the approach using absolute ecosystem service values). Using the median values of the frequency distributions, we calculate the best landscape composition where median landscape production is above a certain % of its maximum value, i.e. function threshold. This is performed for all levels of crop pollination dependence (i.e. for a wide range of crop types) to show the best landscape composition as a function of crop pollination dependence (Fig. 1C). We use median values because they are the most frequent values; besides, median values are robust to non-Gaussian distributions, which are increasingly typical when stochasticity is high. Using any other quantile yields qualitatively similar results (Appendix B).

**Figure 1.**
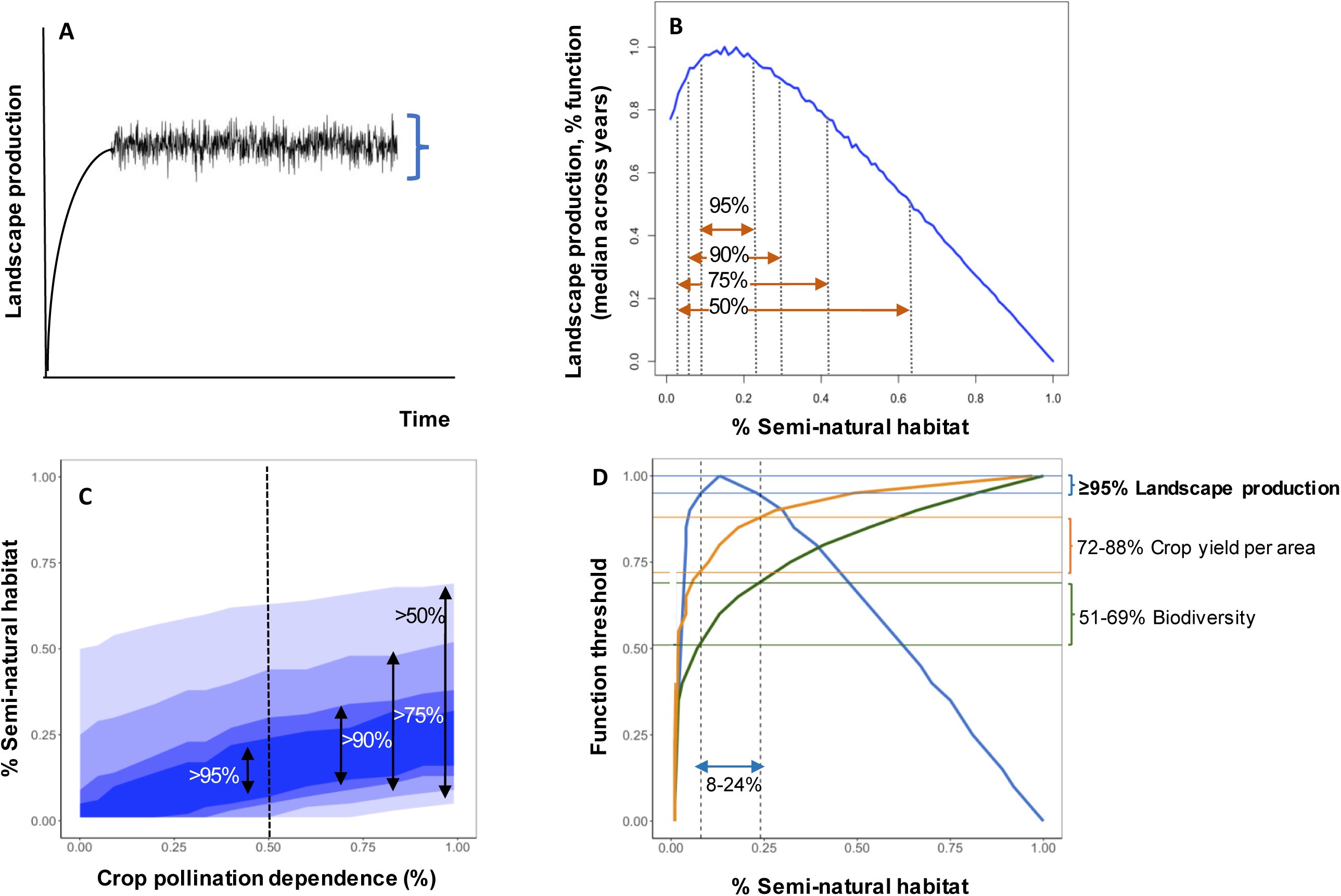
Analytical protocol. We followed a 4-step procedure to estimate the best landscape compositions for each stakeholder’s demand and its consequences on the provision of other ecosystem services. For illustrative purposes, we assume the agricultural unions’ perspective (prioritizes total landscape production). For each fraction of semi-natural habitat, we run model simulations for 1000 time steps (**1A**), which are used to obtain the frequency distribution of landscape production. Once performed for the whole gradient of semi-natural habitat, the distribution of landscape production as a function of semi-natural habitat is obtained (**1B**). This approach enables to explore the full probability distribution of ecosystem service values as well as the possibility of setting thresholds values for services (Carnus et al., 2014). Absolute thresholds for ecosystem services are generally lacking, so we investigate the best landscape compositions that provide ≥95% of landscape production along the range of semi-natural habitat (see Appendix B for an alternative application based on absolute values of ecosystem services). Using the median values of landscape production (blue line), we calculate the best landscape composition (range of % of semi-natural habitat) where median crop yield is above a certain % of its maximum value, i.e. function threshold (here, 95%, 90%, 75% and 50% function thresholds are shown). We complete model simulations for all levels of crop pollination dependence to show the best landscape composition as a function of crop pollination dependence (**1C**). The vertical dashed line represents 50% of pollination dependence of crops (B). The % in (C) correspond to the functional thresholds, and the Y axis shows the range of semi-natural habitat at which the functional thresholds are achieved. More flexible demands (lower function thresholds) imply a wider range of semi-natural habitat values. Finally, (**1D**) shows the range of semi-natural habitat (delimited by dashed vertical lines) that achieves ≥95% of landscape production (agricultural unions’ goal, highlighted in bold). The shadow area represents the % of provision of the other two ecosystem services that corresponds to that range of semi-natural habitat. Figs 1B, D are calculated for 50% of crop pollination dependence, and median values of environmental and demographic stochasticity.

Next, we address the effects of each stakeholder’s demand on the other ecosystem services. For each stakeholder (scenarios 1-3), we set the function threshold of the ecosystem service they demand to 95%, i.e. agricultural unions demand ≥95% of landscape production. The demand established by the function threshold is satisfied within a certain range of semi-natural habitat (Fig. 1D), which is used to determine the corresponding provision of the other ecosystem services (Fig. 1D). For the social average scenario (scenario 4), we apply a *maximin* approach, a common method of performing multi-objective optimization (Solteiro Pires et al., 2005; for general multi-criteria decision analysis, see also Huang et al. 2011, Mendoza and Martins 2006). This method selects the best landscape composition that maximises, within the set of three ecosystem services, the provision of the least provisioned one. In sum, this protocol produces the best landscape compositions for different stakeholders’ demands (scenarios 1-4), crop types (degrees of pollination dependence of crops), and function thresholds, and explores the effects of such demands on other ecosystem services.

Finally, we compare the best landscape compositions obtained by our model with current management of agricultural landscapes. Our search for actual policies targeting specific fractions of semi-natural habitat within agricultural landscapes yielded only one result, the European Union Green Policy, which aims at preserving 5% of semi-natural elements at the farm level (Pe’er et al., 2014). Additionally, we assessed two conservation policies focused on terrestrial ecosystems: the 2020 Aichi Biodiversity Targets (ABT; Global Biodiversity Outlook, 2014) and the Brazilian Forest Code (BFC; Soares-Filho et al., 2014).

## 3. RESULTS

### 3.1. Best landscape compositions for each stakeholder

The best landscape composition depends on stakeholders’ demands, and thus on which ecosystem service is prioritized. In general, the best landscape compositions for individual farmers and conservationists were associated with higher fractions of semi-natural habitat, whereas agricultural unions’ demands were achieved at lower fractions of semi-natural habitat. For intermediate levels of crop pollination dependence and median values of stochasticity, prioritizing individual farmers’ demands yields intermediate-to-high fractions of semi-natural habitat (Fig. 2A). While this favors biodiversity conservation, landscape production is highly variable (1-69%; Figs 2A, 3B). Whereas total production at landscape scale depends on both yields (i.e. related to semi-natural habitat) and crop land, the trade-off between crop yields per area and total production at the landscape scale is explained because yields per unit area increase with more pollinator supply, which in turn increases with semi-natural land. Agricultural unions’ goal to maximize landscape production occurs at low-intermediate fractions of semi-natural habitat (Fig. 2B). This leads to relatively high levels of individual farmers’ demands, but conservationists’ goals may not be fulfilled as biodiversity would remain at intermediate levels (Figs 2B, 3B). More biodiversity remains at higher fractions of semi-natural habitat (Fig. 2C), and it is positively correlated with high levels of crop yield per area, reconciling conservationists’ demands and farmers’ profitability (Figs 2C, 3B). Conversely, landscape production would be low (0-24%), revealing a trade-off between conservationists’ and agricultural unions’ demands. Finally, the social average scenario corresponds to intermediate fractions of semi-natural habitat (39%; Fig. 2D), yielding 79% of the function threshold for all three ecosystem services, which can be considered as indicative of a highly multifunctional agricultural landscape.

**Figure 2.**
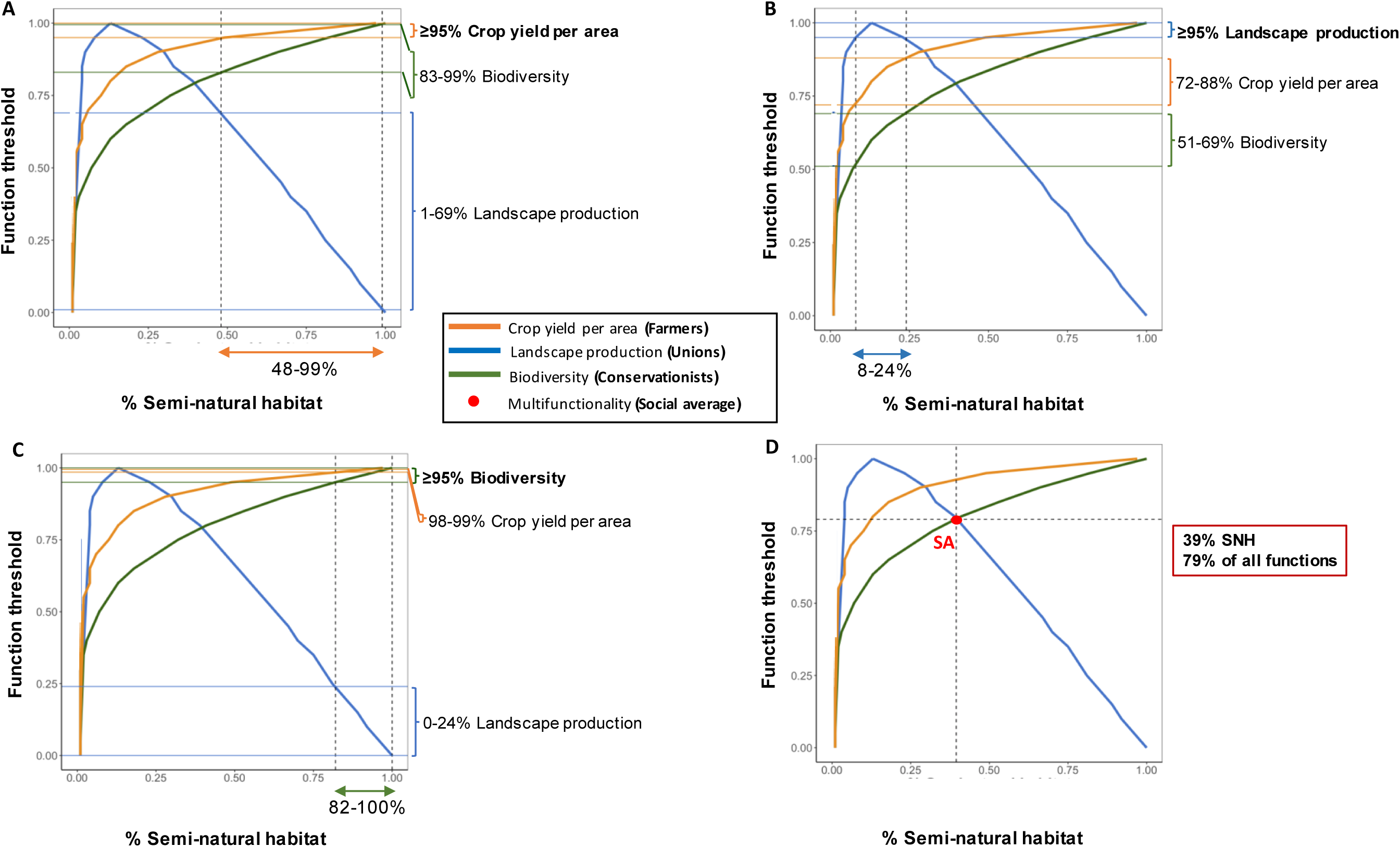
Example of the methodological approach. (**A, B, C**) show the range of semi-natural habitat (delimited by dashed vertical lines) that achieves ≥95% of the stakeholder or ecosystem service prioritized (highlighted in bold in the right-hand side of each plot). The shadow area in (A-C) represent the % of provision of the other two ecosystem services that corresponds to that range of semi-natural habitat. (**D**) The social average scenario is calculated using the *maximin* approach, a common method for multi-objective optimization. This method selects the best landscape composition that maximizes, within the set of three ecosystem services, the provision of the least provisioned one. In this example, the social average scenario (SA) achieves a 79% of all ecosystem services at 39% of semi-natural habitat (red dot). Figs A-D are calculated for 50% of crop pollination dependence and median values of environmental and demographic stochasticity.

**Figure 3.**
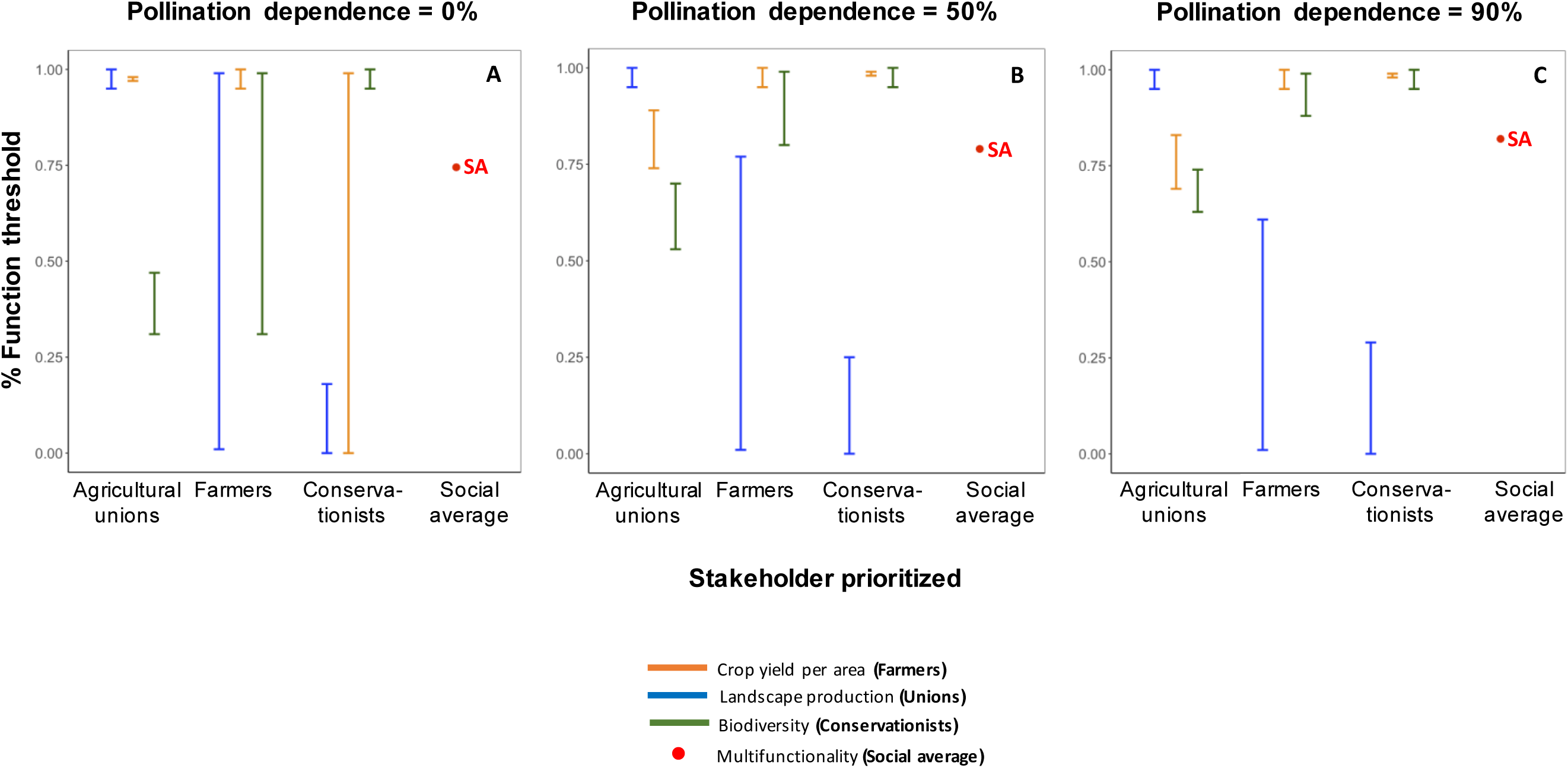
Effects of stakeholders’ demands on the provision of other ecosystem services. In each plot, the stakeholder prioritized is on the X axis (function threshold of its most valued ecosystem service is set to ≥95%), and its effects on the amount of the other two ecosystem services is analyzed. Pollination dependence of crops increases from left to right (A-C). High levels of crop yield per area and biodiversity conservation are compatible, suggesting that individual farmers and conservationists’ demands can be reconciled; this compatibility increases with crop pollination dependence. Landscape production and biodiversity conservation are highly incompatible when agricultural unions’ demands are prioritized. Agricultural unions and individual farmer’s demands can be reconciled when the degree of pollination dependence of crops is low. The social average scenario (SA) yields high provision levels for all three ecosystem services (73-82.5%). Figs 1A-C are calculated for median values of environmental and demographic stochasticity.

### 3.2. Effects of crop type, stochasticity and function thresholds on landscape composition

The best landscape compositions, and the trade-offs and synergies between stakeholders’ demands, strongly depend on three factors: crop pollination dependence, the strength of environmental and demographic stochasticity, and the function threshold of ecosystem services, i.e. the relative amount of an ecosystem service demanded by each stakeholder.

#### 3.2.1 Crop pollination dependence

Higher pollination dependence of crops shifts the best landscape compositions to higher fractions of semi-natural habitat for individual farmers and agricultural unions (Fig. 4, intermediate stochasticity). For increasing levels of pollination dependence, crop yield per area and total landscape production require more pollinators, and hence semi-natural habitat; thus, compatibility between conservationists’ and individual farmers’ demands increases with the level of pollination dependence of crops. Compatibility of union and conservationist demands improves as well given their ranges move closer, yet they do not overlap. Conservationists and agricultural unions’ goals cannot however be reconciled for any type of crop if both stakeholders target ≥95% of their most valued ecosystem service. Only at low levels of crop pollination dependence can agricultural unions and individual farmers coincide in their best landscape composition, but this leads to low biodiversity levels (Fig. 3A). For the social average scenario, the fraction of semi-natural habitat increases with crop pollination dependence, this mainly driven by the dependency of landscape production and crop yield per

**Figure 4.**
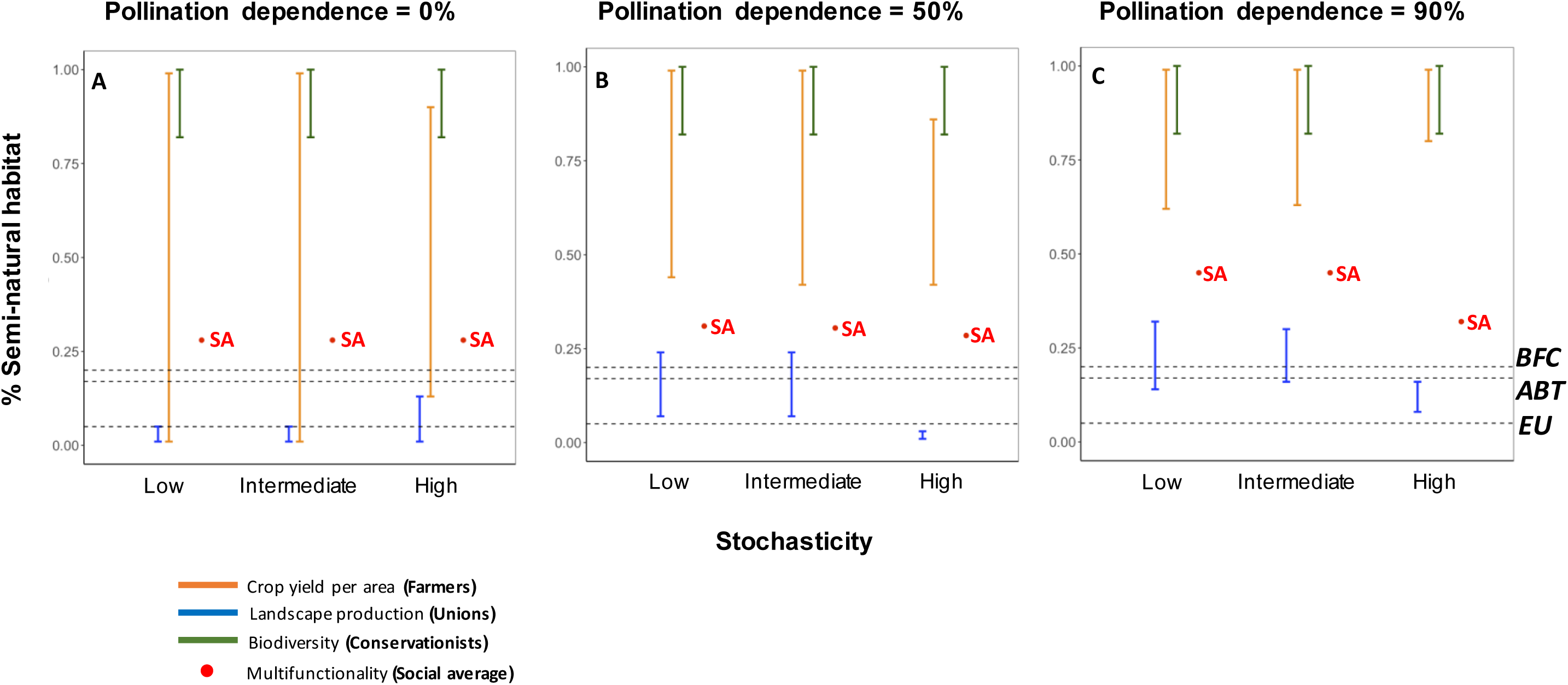
Effects of crop pollination dependence and stochasticity on landscape composition. Function threshold for individual stakeholder’s demand are set to ≥95%. (A-C) Fraction of semi-natural habitat as a function of stochasticity (minimum, median and maximum values for all parameters, both demographic and environmental, are plotted). The effects of stochasticity are more pronounced for high values of stochasticity. Maximum stochasticity shifts the best landscape composition to higher values of semi-natural habitat for landscape production and multifunctionality, and reduces the range width of semi-natural habitat for crop yield per area. The social average scenario (SA) is achieved at higher fractions of semi-natural habitat when the degree of pollination dependence of crops is intermediate and high. The dashed horizontal lines represent the fraction of semi-natural habitat targeted by the EU Green Policy (*EU*), the Aichi Biodiversity Targets (*ABT*) and the Brazilian Forest Code (*BFC*).

#### 3.2.2 Stochasticity

A higher stochasticity increases the fraction of semi-natural habitat that meets individual farmers’ demands, but it decreases the best fraction of semi-natural habitat for agricultural unions, especially for animal-pollinated crops (Figs 4B-C), revealing a trade-off between the two stakeholders. In general, a high stochasticity only slightly reduces the compatibility between farmers and conservationists, but it completely decouples demands of individual farmers and agricultural unions. The latter is due to a large reduction in the fraction of semi-natural habitat to meet agricultural unions’ demands when stochasticity is high together with an increment of the fraction of semi-natural habitat for farmers’ goals. The social average scenario is met at intermediate fractions of semi-natural habitat (30.5, 39 and 45%, for different crop types), but high stochasticity reduces that fraction to lower values (28.5, 29.5 and 31%) because of changes in agricultural unions’ demands. We found a greater effect of environmental (compared to demographic) stochasticity of crops and wild pollinators in determining changes in the best landscape compositions (Fig. C1, Appendix C).

#### 3.2.3 Function threshold

Compatibility among stakeholders’ demands increases with diminishing function thresholds, i.e. the relative amount of an ecosystem service demanded by each stakeholder (Fig. S3). Lower function thresholds expand the range of semi-natural habitat associated with the best landscape composition of each stakeholder. This increases the overlap in the best landscape composition of different stakeholders. Compatibility of stakeholders’ demands with varying function thresholds is higher for intermediate and high levels of crop pollination dependence.

## 4. DISCUSSION

An efficient, sustainable design of agricultural systems is a major challenge of our time, and the integration of various services and stakeholders is crucial to meet that challenge. Our study adds to the large body of literature on how to make management decisions involving multiple objectives, including multi-criteria decision analyses based on multiple valued outcomes (e.g. see reviews by Huang et al., 2011; Mendoza and Martins 2006), and studies addressing the role of ecosystem trade-offs and the role of demand of multiple stakeholders (Landis 2017; Cavender-Bares et al., 2015; Palacios-Agundez et al., 2015; Yahdjian et al., 2015; Burkhard et al., 2012), and highlights the importance of considering the multifaceted nature in the supply of and demand for ecosystem services. Further, our model assesses the potential effects of global change on ecosystem services, thus explicitly including food stability in ecosystem service research (Cruz-Garcia et al., 2016). Our simple case-study model of three ecosystem services and three groups of stakeholders contributes to develop a general understanding of the balance of biodiversity and crop production, and of the various stakeholders, in intensively-managed agricultural systems based on landscape composition. Our results reveal trade-offs and compatibilities between stakeholders’ demands that mirror those observed in the supply side, indicating that prioritization of individual stakeholders’ demands has consequences on the other services, and on ecosystem multifunctionality. Such trade-offs strongly depend on factors seldom considered by management policies, including the degree of pollination dependence of crops and the strength of environmental and demographic stochasticity.

The best landscape compositions, measured as the fraction of semi-natural habitat within the agricultural landscape, differ among stakeholders. In general, individual farmers’ and conservationists’ demands are associated with higher fractions of semi-natural habitat, whereas agricultural unions’ demands are achieved at lower fractions of semi-natural habitat. This trade-off in the ecosystem service demand between agricultural unions, on one hand, and individual farmers and conservationists, on the other hand, implies that management focusing on single stakeholders will invariably reduce multifunctionality in agricultural systems, at least for the three services considered in this study. But the best landscape composition is not a static figure; rather, it depends on the same factors that determine trade-offs in the provision of ecosystem services in agricultural landscapes (Montoya et al., 2018).

Crop type, defined here as the degree to which a crop depends on animal pollination, is a main driver of the best landscape composition for both individual farmers and agricultural unions. A higher pollination dependence of crops generally shifts the best landscape composition to larger fractions of semi-natural habitat, irrespective of the stakeholder considered. Yet, to our knowledge this shift is not taken into account by actual policies. For low levels of pollination dependence of crops and crops independent of animal pollination, prioritizing agricultural unions’ demands increases landscape production, but reduces farmers’ benefits and harms biodiversity significantly. However, with increasing dependence of crops on animal pollination the best landscape composition involves larger fractions of semi-natural habitat, which results in better biodiversity conservation combined with farmers’ profitability, but does not satisfy agricultural unions’ demands. Because agriculture worldwide is becoming more pollinator-dependent over time (Breeze et al., 2014; Aizen et al., 2009), changes in the best landscape compositions driven by crop pollination dependence are highly relevant in the global context.

Current global change enhances the inter-annual variance of several climate variables, i.e. environmental stochasticity (Giorgi et al., 2001; Li and Xian, 2003; Saltz et al., 2006), as shown by the increasing frequency of extreme climatic events such as floods, heat waves and droughts (Fischer et al., 2016; Woodward et al., 2016; Craven et al., 2016). These events may in turn increase environmental stochasticity of crops and wild pollinators, and there is evidence that climate trends are partly responsible for observed increases in yield variability (Iizumi and Ramankutti, 2016; Osborne and Wheeler, 2016), and influence the delivery of ecosystem services (Shoyama & Yamagata 2014; Ding et al., 2016; Martinez-Harms et al., 2017). Our results suggest that increasing stochasticity has contrasting effects on the best landscape composition for different stakeholders. Individual farmers’ demands are generally met at higher fractions of semi-natural habitat when stochasticity is high, whereas the opposite holds true for agricultural unions. Such differences are driven by the different effects of stochasticity on the provision of ecosystem services. More specifically, high stochasticity changes the relationship between ecosystem services and semi-natural habitat: on one hand, the unimodal relationship of landscape production reported elsewhere (Braat and ten Brink, 2008; Montoya et al., 2018) becomes monotonically decreasing for high stochasticity (Appendix C, Fig. C1), and this shifts maximum landscape production to lower fractions of semi-natural habitat; on the other hand, the saturating relationship of crop yield per area becomes flat when stochasticity is high (Appendix C, Fig. C2), thus constraining the range of semi-natural habitat that maximizes farmers’ profitability. Agricultural management should thus consider global change predictions, as changes in stochasticity directly influence the best landscape compositions that maximize stakeholders’ demands.

The demands of individual farmers and conservationists generally align at higher fractions of semi-natural habitat. Therefore, the farmers’ best interest is that, despite how much farmed is their land, the landscape around their cultivated land is not farmed. This is consistent with empirical research in grassland ecosystems showing that higher levels of biodiversity are beneficial for landowners (Binder et al., 2018), but contrasts with recent trends in modern agriculture, where individual farmers tend to increase crop land at the expense of semi-natural habitat, and thus biodiversity. We identify various reasons to explain the discrepancy between the individual farmers’ demand defined in our study and their ‘actual’ demand. First, crops whose production does not depend on wild pollination do not require semi-natural habitat; in this case, farmers tend to expand crop area. Second, wild pollinators might suffer from the tragedy of the commons, where wild pollinators are the commons and farmers deplete semi-natural habitat through their collective action. Finally, the importance of wild pollinators inhabiting non-crop areas for crop production – pollination of crop plants – might be underestimated.

Our results suggest that a reduction in the amount of an ecosystem service that stakeholders are willing to accept (i.e. function thresholds) is a necessary condition to reconcile various stakeholders’ demands (Figs 2D, S3). This is because lower function thresholds expand the range of semi-natural habitat, increasing the overlap in the best landscape composition for different stakeholders. Indeed, this idea underpins the social average scenario, which suggests that overall performance of agricultural landscapes can be improved by combining multiple demands for ecosystem services, as opposed to a traditional focus constrained by provisioning services, mainly total landscape production (Lovell and Johnston, 2009; Jordan and Warner, 2010). Our results suggest that prioritizing multifunctional landscapes achieves relatively high levels of the three ecosystem services analysed and satisfies stakeholders’ demands for intermediate amounts of semi-natural habitat. Further, the social average scenario is more robust to changes in crop type and stochasticity (Figures 3, 4, S1). Therefore, management of agricultural systems for multifunctionality may better align with the increasing consensus supporting the need for agricultural landscapes to simultaneously provide ecosystem services that guarantee food security, livelihood opportunities, and biodiversity conservation (i.e. ecosystem service multifunctionality, Manning et al., 2018).

Our findings suggest that, unless the amount of an ecosystem service that stakeholders are willing to accept is reduced, the proportion of semi-natural habitat that conservation policies aim to protect lies below the best landscape composition in intensively-managed agricultural systems, especially for high levels of crop pollination dependence. The EU Green Policy succeeds in achieving high levels of landscape production (agricultural unions’ demand) of crops that do not depend much on animal pollination, although this leads to low biodiversity levels (Figs 4, S1, S3). Similar conclusions can be drawn for the Aichi Biodiversity Targets and the Brazilian Forest Code, although they target higher fractions of semi-natural habitat (17% and 20%, respectively). Therefore, the regional and global-scale policies analyzed here satisfy agricultural unions’ demands in one scenario only, that is, when crops depend little (or not at all) on animal pollination, although it leads to low biodiversity levels. By and large, preserving 5-20% of semi-natural habitat lies below the best landscape composition for biodiversity and crop production in intensively-managed agricultural systems for any other combination of stakeholders’ demands considered here, crop type, or stochasticity. Such policies also fail to meet the social average scenario. Under the current trends of increasing pollinator-dependence of agriculture and global change, such targets seem too low to either meet stakeholders’ demands or achieve adequate, sustainable levels of multifunctionality in agricultural systems (social average scenario).

Our approach has several limitations. For example, our results refer to intensively-managed agricultural systems, where crop land does not harbor important biodiversity levels; in non-intensive agricultural systems, however, the best landscape compositions may not be necessarily similar (Clough et al., 2011). Also, we have used landscape composition as our key variable, but other metrics to compare management scenarios exist (e.g. agrochemical inputs), and can be used complementarily. Besides, our model does not consider the effects of the spatial configuration of semi-natural habitat on the best landscape composition, which is expected to determine ecosystem service flows between crop land and fragments of semi-natural habitat (Garibaldi et al., 2011b; Mitchell et al., 2015). Honeybee colonies are frequently used to substitute wild pollinator communities, and our results would be affected if we consider managed honeybees as they do not depend on the availability of semi-natural habitat. However, we purposely did not take managed honeybees into account because it has been argued that the pollination services of wild pollinators cannot be compensated by managed bees due to the fact that pollinator-dependent crop land grows more rapidly than the stock of honeybee colonies (Aizen et al., 2009), wild insects usually pollinate crops more efficiently than honeybees (Garibaldi et al., 2013), and honeybees may depress wild pollinator densities (Lindström et al., 2016). Finally, we have illustrated our approach using three ecosystem services and three groups of stakeholders as case studies; however, agricultural landscapes include a greater variety of ecosystem services (water quality, pest control, flood mitigation, as well as recreational and aesthetic services) and stakeholders whose demands may partially overlap (e.g. farmers may not only favor crop yield per area, but also a minimum amount of crop land). Despite these limitations, our approach is very useful to study how biodiversity and crop production, and the stakeholder demands associated to them, respond to crop type (degree of pollination dependence), environmental and demographic stochasticity, and functional thresholds, based on landscape composition.

### 4.1. Conclusions

The future of intensive farming systems in the context of global change is a key component of the 2017 UN Sustainable Development Goals. Our analysis of three ecosystem services related to biodiversity and crop production, and three groups of stakeholders, shows that the best landscape composition differs among stakeholders, and that current policies should start to consider factors such as crop type, stochasticity, and the amount of an ecosystem service that stakeholders are willing to accept, as they can strongly influence these best landscape compositions for different stakeholders. Management for social average, or multifunctionality scenario, may be a better option for food security, livelihood opportunities, and biodiversity conservation, thus meeting various stakeholders’ demands. Worldwide trends in agriculture (more pollinator-dependent over time) and global change (associated with the strength of environmental and demographic stochasticity) calls for innovative, integrative perspectives in agricultural management that consider a variety of stakeholders and ecosystem services.

## Supporting information

## Author’s contributions

DM, ML and SG conceived the original idea and designed the research. DM, ML and CdM designed the analytical protocol. DM performed the analysis. All authors contributed to interpretation of results. DM wrote the first draft of the manuscript. All authors critically reviewed drafts and have approved the final version.

## Acknowledgements

DM was funded by the EU and INRA in the framework of the Marie-Curie FP7 COFUND People Program, through the award of an AgreenSkills/AgreenSkills+ fellowship. This work was supported by the TULIP Laboratory of Excellence (ANR-10-LABX-41), ANR AGROBIOSE (ANR-13-AGRO-0001), ERANET ECODEAL, and BIOSTASES Advanced Grant, funded by the European Research Council under the European Union’s Horizon 2020 research and innovation program (grant agreement No 666971). We thank Bart Haegeman and the Centre for Biodiversity Theory and Modelling members for helpful discussions.

## Data accessibility

Upon acceptance of the manuscript, we agree to archive any result from simulations in an appropriate public repository.

