## Supplementary material for "Reconciling biodiversity conservation, food production and farmers’ demand"

### **Appendix A**. Model equations and parameter values.

The three components of our model (pollinators, 'wild' plants, and crop yield) are represented by the following equations:

$$\frac{dP}{dt}=r_P(t)P(t)(1-\frac{P(t)}{k_P\omega_{sn}A})+\sigma_P^e u_P^e(t)P(t)+\frac{\sigma_P^d u_P^d(t)}{\sqrt{P(t)}}P(t)$$ Eq. (A.1)

$$\frac{dW}{dt}=r_W(t)W(t)(1-\frac{W(t)}{k_W\omega_{sn}A})+\sigma_W^e u_W^e(t)W(t)+\frac{\sigma_W^d u_W^d(t)}{\sqrt{W(t)}}W(t)$$ Eq. (A.2)

$$C(t)=(1-\omega_{sn})A[Z_C+\frac{\alpha_C[P(t)/A]}{\beta_C+[P(t)/A]}](1+\sigma_C^e u_C^e(t))$$ Eq. (A.3)

where $P$ and $W$ represent the maximum yearly biomass of pollinators and 'wild' plants, respectively. $P$ does not take managed honeybees into account as they do not depend on the availability of semi-natural habitat, and they pollinate less efficiently compared to non-managed pollinators (Garibaldi et al., 2013). The model does not consider within-year dynamics. $C(t)$ is the amount of crop biomass produced in year $t$, i.e. annual crop yield. $C(t)$ is not represented by a differential equation because crops are harvested and their dynamics do not depend on the previous state. Conversely, pollinators and wild plants are not managed and their actual values depend on previous states. $k_P$ and $k_W$ are the carrying capacities of pollinators and 'wild' plants, respectively, per unit area; $A$ is the total landscape area (crop land and semi-natural habitat); $\omega_{sn}$ is the proportion of semi-natural habitat within the agricultural landscape ($[1-\omega_{sn}] * A$ is total crop or agricultural area).

In the first two equations, $r_P(t)$ and $r_W(t)$ are the pollinators' and 'wild' plants' per capita growth rates, and are defined as:

$$r_P(t)=c_P\frac{\alpha_P(\phi_W W(t)+\phi_C C(t))}{\beta_P+\phi_W W(t)+\phi_C C(t)}$$ Eq. (A.4)

$$r_W(t)=c_W\frac{\alpha_W(P(t)/A)}{\beta_W+(P(t)/A)}$$ Eq. (A.5)

Pollinators are assumed to be generalist central-place foragers that feed on both 'wild' plants and crops (Kleijn et al., 2015). The model can apply to any spatial extent provided that the pollinators, and hence the fragments of semi-natural habitat that host them, are distributed in such a way that pollinators have access to the whole landscape. Therefore, the spatial extent can vary from roughly 10 ha (typical arable field in Europe) to any larger scale provided pollinators are not aggregated in a small part of the landscape. We assume that plant and pollinator uptake of resources follows a saturating, type II functional response, where $\alpha_P$ and $\alpha_W$ are the maximum growth rates; $\beta_P$ and $\beta_W$ are half-saturation constants; and $c_P$ and $c_W$ are the conversion rates of pollinators and 'wild' plants, respectively, that translate the functional responses into numerical ones. For simplicity, we set conversion rates equal to unity. The pollination-dependent part of crop yield is also assumed to follow a type II functional response, where $\alpha_C$ is the maximum crop yield derived from pollination, $\beta_C$ is the half-saturation constant of crops, and $\Phi_W$ and $\Phi_C$ are constants that convert fluxes of 'wild' plants and crops, respectively, to pollinator biomass. We use $\Phi_W = \Phi_C = 1$ for simplicity; to allow differences in resource quality of different crop types, we also made $\Phi_C$ dependent on crop pollination dependence (see below). The use of saturating functional responses is widely supported and it is consistent with several biological examples (Thebault & Fontaine, 2010; Holland et al., 2013; Holland, 2015).

Environmental stochasticity (e.g. rainfall variability, variation in temperature) is included through the terms $\sigma^e u^e(t)$, where $(\sigma^e)^2$ is the environmental variance of either pollinators ($(\sigma_P{}^e)^2$), 'wild' plants ($(\sigma_W{}^e)^2$) or crops ($(\sigma_C{}^e)^2$), and $u^e(t)$ are random functions with zero mean and standardized variance, that can be correlated through time. Demographic stochasticity ($\sigma^d u^d(t)$) emerges from stochastic variation in individuals' births and deaths. Crops are sown at high densities, and thus we assume demographic stochasticity is prevented in crops, and only affects pollinators and 'wild' plants. Demographic stochasticity is included in the form of the first-order normal approximation commonly used in stochastic population dynamics (Lande et al., 2003), where $(\sigma^d)^2$ is the demographic variance of either pollinators ($(\sigma_P{}^d)^2$) or 'wild' plants ($(\sigma_W{}^d)^2$), and $u^d(t)$ are independent random functions with zero mean and standardized variance.

Crops differ greatly in the degree to which animal pollination contributes to yield, from pollinator-independent crops, such as obligate wind- or self-pollinated species (e.g. cereals), to fully animal-pollinated species (e.g. fruit trees, oilseed rape). Within animal-pollinated species, crops differ in their level of dependence on pollination (Klein et al., 2007). In our model, $Z_C$ represents the part of crop yield that is independent of animal pollination and $\alpha_C$ is the crop yield derived from pollination, and therefore we can estimate crop pollination dependence (%) as $\alpha_C / (\alpha_C + Z_C)$. If $Z_C = 0$ ($\alpha_C > 0$), crop yield depends entirely on animal pollination; conversely, animal pollination-independent crops are defined by $\alpha_C = 0$ ($Z_C > 0$). Most fruit and seed crops lie between these two extremes ($Z_C > 0$, $\alpha_C > 0$).

We use the species-area relationship (SAR) to estimate changes in pollinator biodiversity as a function of semi-natural area. Despite SAR is usually stronger at spatial scales larger than that of arable fields, where we might observe more variation around the average biodiversity values, it captures the expected mean biodiversity at the scale of an arable field in Europe. We estimated SAR using the conventional power law function ($S=c\,[\omega_{sn}\,A]^z$, where $S$ = number of species, $c$ is a constant of proportionality). Theoretical models and field data from a wide range of plant and animal taxa show that the slope, $z$, of the logarithm of species richness against the logarithm of area is roughly constant, with $z \approx 0.25$ (Crawley and Harral, 2001).

We estimated parameter values using empirical information and commonly-assigned values found in the literature (McCann et al., 2005; Thompson et al., 2006; Leroux and Loreau, 2008; Holland and DeAngelis, 2010; Thebault and Fontaine, 2010; Morales, 2011; Holland et al., 2013; Encinas-Viso, 2014; Gounand et al., 2014). For example, to determine the carrying capacity of pollinators ($k_P$), we used empirical data on average numbers of individuals and body mass of wild pollinators (Bommarco et al., 2012; Rollin et al., 2013; Holzschuh et al., 2016). For wild plants, we used empirical observations to inform their carrying capacities ($k_W$) (Craven et al., 2016). Also, there is information on independent crop yield that was used to determine $Z_C$ (e.g. http://data.worldbank.org/). Environmental and demographic stochasticity values were obtained from experimental studies (de Mazancourt et al., 2013). We allowed variation in $\alpha_C$ and $\beta_C$ to investigate changes in ecosystem services across the amount of semi-natural habitat ($\omega_{sn}$), and the degree of crop pollination dependence ($Z_C/\alpha_C$). Sensitivity analysis shows our results are robust to changes in parameter values (Montoya et al., 2018).

**Table A.1.** Parameters and variables of the model

| Parameters & Variables | Definition | Dimensions |
|---|---|---|
| Parameters | | |
| $\alpha_P$ | Maximum growth rate of pollinators | $time^{-1}$ |
| $\alpha_W$ | Maximum growth rate of semi-natural plants | $time^{-1}$ |
| $\alpha_C$ | Maximum crop yield derived from pollinator interactions | $mass \cdot area^{-1}$ |
| $\beta_P$ | Half-saturation constant of pollinators | mass |
| $\beta_W$ | Half-saturation constant of 'wild' plants | $mass \cdot area^{-1}$ |
| $\beta_C$ | Half-saturation constant of crop plants to pollinators | $mass \cdot area^{-1}$ |
| $k_P$ | Carrying capacity of pollinators per unit area | $mass \cdot area^{-1}$ |
| $k_W$ | Carrying capacity of semi-natural plants per unit area | $mass \cdot area^{-1}$ |
| $A$ | Total landscape area | area |
| $\omega_{sn}$ | Proportion of semi-natural habitat | dimensionless |
| $Z_C$ | Crop yield independent of pollinators | $mass \cdot area^{-1}$ |
| $c_W$ | Conversion rate of 'wild' plants (from functional to numerical response) | dimensionless |
| $c_P$ | Conversion rate of pollinators (from functional to numerical response) | dimensionless |
| $\Phi_W$ | Weighting factor for 'wild' plants (flux to stock) | dimensionless |
| $\Phi_C$ | Weighting factor for crop plants (flux to stock) | dimensionless |
| $r_P$ | Intrinsic growth rate of pollinators | $time^{-1}$ |
| $r_W$ | Intrinsic growth rate of 'wild' plants | $time^{-1}$ |
| $\sigma^e{}_P$ | Environmental standard deviation of pollinators | $time^{-1/2}$ |
| $\sigma^e{}_W$ | Environmental standard deviation of 'wild' plants | $time^{-1/2}$ |
| $\sigma^e{}_C$ | Environmental standard deviation of crop production | dimensionless |
| $\sigma^d{}_P$ | Demographic standard deviation of pollinators | $mass^{1/2} \cdot time^{-1/2}$ |
| $\sigma^d{}_W$ | Demographic standard deviation of semi-natural plants | $mass^{1/2} \cdot time^{-1/2}$ |
| $u^e{}_P$, $u^d{}_P$, $u^e{}_W$, $u^d{}_W$, $u^e{}_C$, $u^d{}_C$ | White noise signals with zero mean and standardized variance. $u^e$ = environmental, $u^d$ = demographic $P$ = pollinators; $W$ = 'wild' plants; $C$ = crop plants | dimensionless |
| Variables | | |

| | | |
|---|---|---|
| *C (t)* | Biomass of crop plants (crop yield) | mass |
| *W(t)* | Biomass of semi-natural or 'wild' plants | mass |
| *P(t)* | Biomass of pollinators | mass |
