## Supplementary material for "Reconciling biodiversity conservation, food production and farmers’ demand"

### **Appendix B**. Applications of our methodological approach.

This appendix shows an example of an additional application of our analytical approach. As described in the main text, we follow a several-step process. First, for each fraction of semi-natural habitat, we run model simulations to obtain the frequency distribution for each ecosystem service after 1000 time steps (hereafter, total landscape production is used as example ecosystem service; Figure B.1A-B below). This is performed for the whole range of semi-natural habitat (0-100%) to obtain the frequency distribution of total landscape production as a function of semi-natural habitat (Figure B.1C below).

***Figure B.1***. *Methodological approach. The plots are calculated for crops with a degree of animal pollination of 50%, and median values of environmental and demographic stochasticity.*

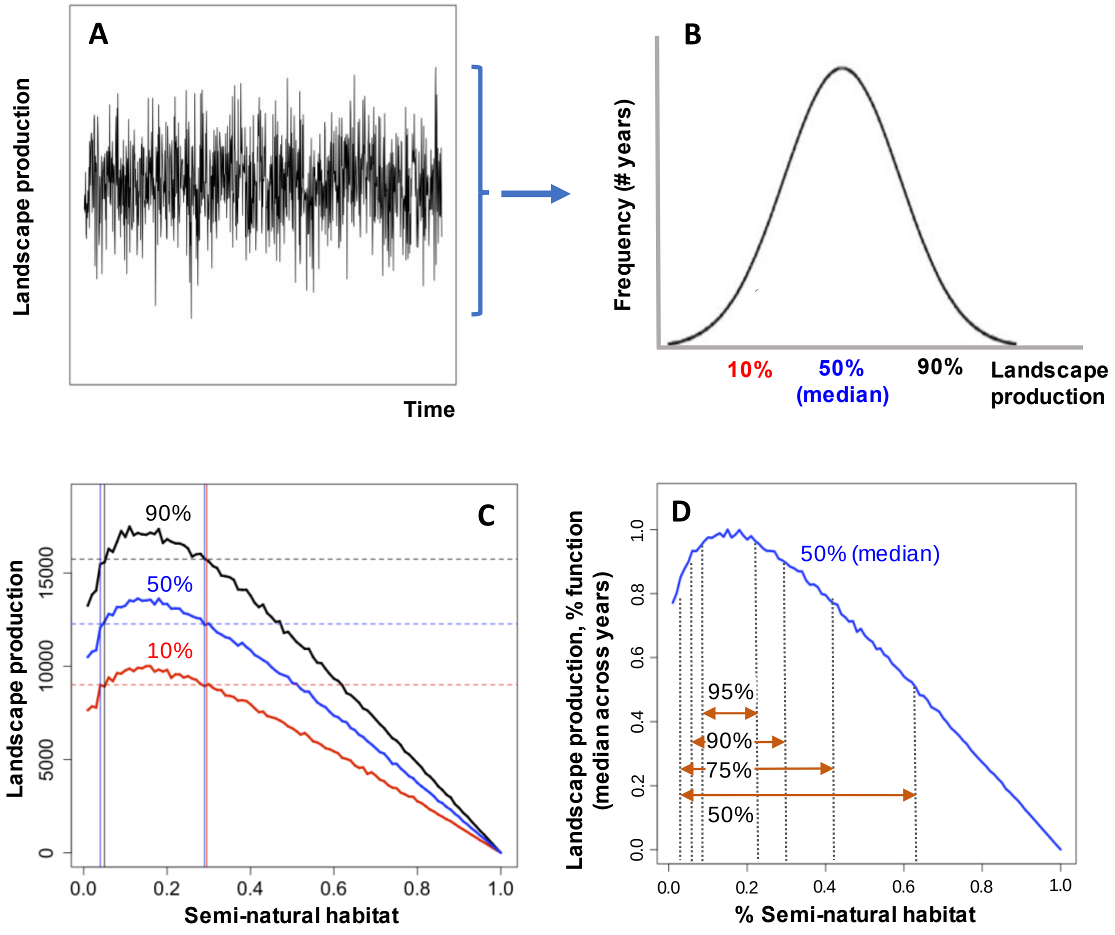


The curves in figure B.1C correspond to three different quantiles (10%, 50%, 90%) of the frequency distribution shown in figure B.1B. An alternative application to the approach we use in the main text consists in defining certain absolute thresholds for ecosystem service values, and looking at the probability of being above those thresholds with time, i.e. frequency of years that landscape production is above a given absolute value. The horizontal dashed lines in figure B.1C represent three examples of such absolute values: 9000 kg (red), 12000 kg (blue), and 16000 kg (black). For the particular case of crops with a degree of animal pollination of 50%, and median values of environmental and demographic stochasticity, those absolute thresholds are guaranteed for >90%, >50%, and >10% of years (Figure B.1C). This alternative application of our approach would be very relevant as maximum values of ecosystem services are often unknown. Unfortunately, absolute thresholds for ecosystem services are generally lacking, so in the main text we look at optimal landscape compositions that provide a given percentage of the maximum ecosystem value (i.e. function threshold) along the range of semi-natural habitat (Figure B.1D). To do this, we use median values because they are the most frequent values, and due to their robustness to non-Gaussian distributions, which are increasingly typical when stochasticity is high. Yet, using any other quantile yields qualitatively similar results because all quantile curves have similar shapes (approximately same optimum semi-natural habitat and same slopes), which leads to similar ranges of semi-natural habitat for the different quantile curves, as shown by the vertical lines in figure B.1C. In conclusion, the strength of this approach lies in the ability to explore the full probability distribution of ecosystem service values as well as the possibility of setting thresholds values for services.
