## Supplementary material for "Reconciling biodiversity conservation, food production and farmers’ demand"

### **Appendix C**. Effects of stochasticity on ecosystem services

This appendix shows the effects of individual stochasticity parameter on ecosystem services. For each of the plots below, each individual parameter (environmental or demographic) is set to its maximum empirical value while the rest are kept to their median values.

***Figure C.1**. Changes in individual stochasticity parameters and their effects on landscape production (Env = environmental stochasticity; Dem = demographic stochasticity; P = pollinators; W = Wild plants; C = crops). The dashed vertical lines represent the range of semi-natural habitat where ≥95% of the maximum landscape production is obtained. Landscape production for median values of all stochasticity parameters is shown for comparison purposes.*

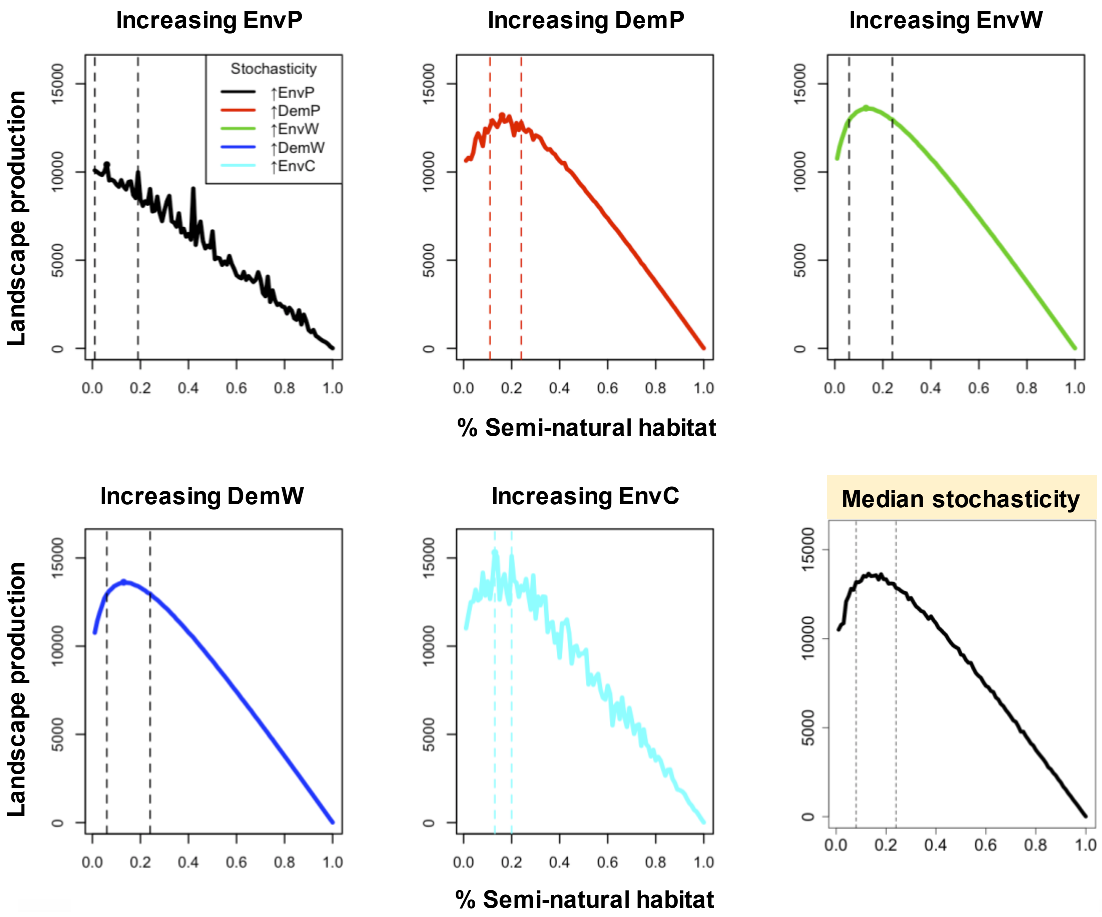


Figure C.1 shows that environmental stochasticity of pollinators (EnvP) increases variability of landscape production, but, most importantly, it erodes the unimodal relationship of landscape production as a function of semi-natural habitat reported in previous studies (Braat & ten Brink, 2008; Montoya et al., 2018). Demographic stochasticity of pollinators (DemP) increases variability only at low fractions of semi-natural habitat. Increasing stochasticity (either environmental or demographic; EnvW, DemW) of wild plants does not affect landscape production. Environmental stochasticity of crops (EnvC) increases variability, but does not change the overall shape of the relationships. Therefore, the larger impact on landscape production is due to increasing environmental stochasticity of pollinators and crops, which influences the optimal landscape composition for landscape production.

***Figure C.2***. *Changes in individual stochasticity parameters and their effects on crop yield per unit area (Env = environmental stochasticity; Dem = demographic stochasticity; P = pollinators; W = Wild plants; C = crops). The dashed vertical lines represent the range of semi-natural habitat where ≥95% of the maximum crop yield per unit area is obtained. Crop yield per are for median values of all stochasticity parameters is shown for comparison purposes.*

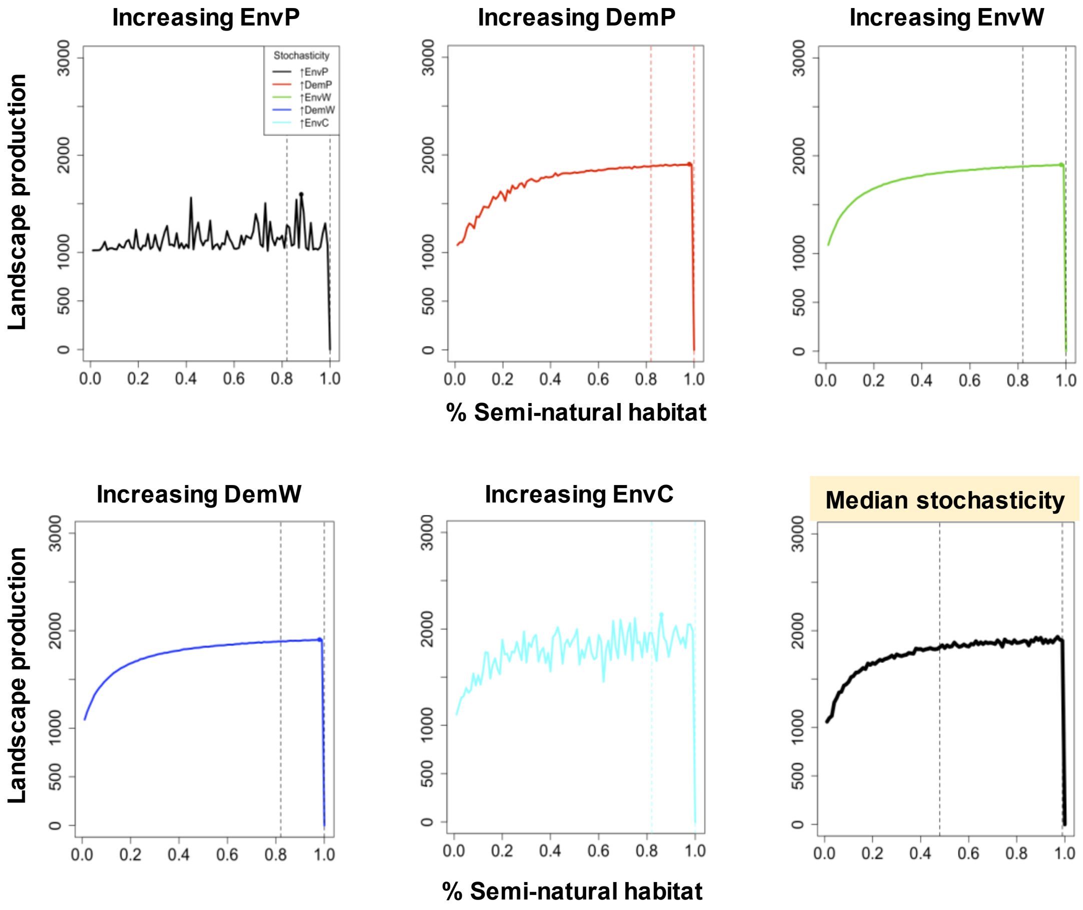


Figure C.2 also shows a higher impact of increasing environmental stochasticity of pollinators and crops on crop yield per unit area, and therefore on the optimal landscape composition: whereas envC increases fluctuations of crop yield per area as a function of semi-natural habitat, high envP also flattens out the relationship between crop yield per area and semi-natural habitat (Montoya et al., 2018), influencing the range of semi-natural habitat that maximizes crop yield per area. Increasing stochasticity (either environmental or demographic) of wild plants has a negligible effect on crop yield per area, whereas high demographic stochasticity of pollinators only affect crop yield per area at low fractions of semi-natural habitat.
